# Prior *Bombyx mori* latent virus exposure alters subsequent BmNPV infection in the silkworm, *B. mori*

**DOI:** 10.64898/2026.10.14.758696

**Authors:** Nura Abubakar, Hiroaki Mon, Tsuguru Fujii, Toshiaki Fujimoto, Takahiro Kusakabe, Masato Hino

## Abstract

*Bombyx mori* latent virus (BmLV) is a persistent RNA virus frequently detected in silkworm-derived cell lines and has been reported to interact with *B. mori* nucleopolyhedrovirus (BmNPV). However, whether previous BmLV exposure alters subsequent BmNPV infection in silkworm larvae remains unclear. Here, we examined how BmLV exposure history affects subsequent BmNPV infection in cultured cells and fifth-instar *B. mori* larvae. Persistent BmLV infection increased BmNPV-mediated reporter protein expression and BmNPV genome abundance in cultured cells, whereas short-term BmLV exposures, including simultaneous exposure to BmLV and BmNPV, did not reproduce these effects. In contrast, 24 h BmLV pre-exposure before BmNPV challenge increased BmNPV-mediated reporter protein expression and BmNPV genome abundance in f38 and n22 silkworm larvae. BmLV antigenome RNA was detected in multiple larval tissues after BmLV exposure, indicating BmLV replication during the pre-exposure period. Notably, following subsequent BmNPV infection, BmLV-pre-exposed larvae showed reduced induction of *Dicer-2*, *R2D2*, and stimulator of interferon genes (*STING*) in the fat body. These findings indicate that prior BmLV exposure enhances subsequent BmNPV infection and is associated with attenuated induction of host antiviral genes. This study expands our understanding of how sequential viral infections shape host–virus interactions and suggests that BmLV exposure history may influence baculovirus-based recombinant protein expression.

**Highlights:**

- BmLV pre-exposure increases BmNPV propagation and reporter protein expression
- BmLV pre-exposure attenuates induction of selected antiviral genes
- BmLV replicates in multiple larval tissues upon experimental exposure.
- Persistent BmLV exposure increases subsequent BmNPV infection in silkworm cells.
- Short-term BmLV exposure or coinfection does not mimic persistent BmLV infection

## Introduction

Animals are frequently exposed to multiple pathogens during their lifetime, and infection history is known to influence the outcome of subsequent infections. In insects, previous viral exposure can alter the outcome of subsequent infection, ranging from enhanced resistance after prior exposure (Tidbury et al., 2011) to increased susceptibility under some exposure conditions (Wilson et al., 2021), while other studies have found no detectable effect (Longdon et al., 2013). In the lepidopteran *Plodia interpunctella*, for example, prior exposure to a low dose of granulosis virus reduced susceptibility to a subsequent challenge, demonstrating that viral infection history can influence antiviral defense within a generation (Tidbury et al., 2011). Whether exposure to a persistent RNA virus can similarly alter susceptibility to a subsequent DNA virus infection, however, remains poorly understood in lepidopteran insects

*Bombyx mori* latent virus (BmLV), formerly described as *B. mori* macula-like virus, provides an experimental system to examine this question. BmLV is a positive-sense single-stranded RNA virus originally identified in the *B. mori* ovary-derived BmN-4 cell line, where it establishes persistent, high-level infection without strong cytopathic effects (Katsuma et al., 2005; Iwanaga et al., 2012). BmLV infection is accompanied by antiviral RNA interference (RNAi) responses, including the production of virus-derived small interfering RNAs (siRNAs) and PIWI-interacting RNA (piRNA)-sized RNAs. Genetic and molecular studies indicate that *Ago2*-and PIWI-associated pathways contribute to controlling BmLV accumulation in persistently infected cells (Katsuma et al., 2018; Katsuma et al., 2021a). Recent work further suggests that BmLV-derived piRNAs can arise through non-canonical processing of abundant viral transcripts rather than a conventional ping-pong pathway (Verdonckt et al., 2026). Thus, persistent BmLV infection provides a sustained interaction between viral RNA and host antiviral pathways, although whether this infection alters the response to a subsequent virus remains unclear.

The biology of BmLV differs between cultured cells and intact silkworms. BmLV establishes persistent infection in silkworm-derived cell lines, where viral RNA is readily detected. In contrast, following experimental exposure of *B. mori* Kinshu Showa larvae, BmLV RNA has been detected in the fat body, midgut, and gonads, with only low levels of viral RNA detected in the ovary (Katsuma et al., 2005; Iwanaga et al., 2012; Iwanaga, 2019; Tsukui et al., 2019; Feng et al., 2021). BmLV-derived small RNAs are also detectable following experimental exposure, although at low abundance, and RNA silencing alone does not appear to explain the resistance of silkworm ovaries to BmLV (Katsuma et al., 2021b).

*B. mori* nucleopolyhedrovirus (BmNPV) is a DNA baculovirus and an important pathogen of *B. mori* (Harrison et al., 2018). In silkworm, BmNPV infection is controlled by multiple host antiviral mechanisms, including RNAi and the *STING* (stimulator of interferon genes)-dependent signaling and other innate immune and cellular defense pathways (Jiang, 2021; Hua et al., 2018; Nagamine and Sako, 2016). RNAi limits viral RNA through small-RNA-guided targeting and degradation, whereas *STING*-dependent signaling contributes to antiviral defense by promoting activation and nuclear translocation of the nuclear factor kappa B (NF-κB)-related transcription factor *BmRelish* following BmNPV infection (Hua et al., 2018; Wei et al., 2025).

Previous studies have also demonstrated interactions between BmLV and BmNPV. Persistent BmLV infection caused a modest delay in BmNPV propagation in cultured cells, whereas simultaneous inoculation of the two viruses into larvae increased BmLV RNA accumulation. Infectious BmLV particles were also detected within BmNPV occlusion bodies produced in BmLV-positive cells (Tsukui et al., 2019). However, it remains unknown whether prior BmLV exposure, including an exposure that produces no overt disease phenotype in larvae, can alter BmNPV propagation and antiviral responses.

In this study, we examined how BmLV exposure influences subsequent BmNPV infection in cultured cells and *B. mori* larvae. We compared BmLV pre-exposure with BmLV coinfection and examined whether prior exposure altered BmNPV genome accumulation, reporter protein expression, and antiviral gene responses.

## Results

### 3.1 Persistent BmLV infection, but not short-term exposure, enhances BmNPV abundance and BmNPV-mediated reporter expression in cultured cells

BmLV is commonly maintained as a persistent infection in *B. mori*-derived cell lines, and previous studies have shown that persistent BmLV infection can affect BmNPV propagation in cultured cells (Tsukui et al., 2019). We therefore examined whether this effect was associated specifically with persistent infection or could also be reproduced by short-term BmLV exposure using BmMAX7050np cells. BmLV-free cells, persistently BmLV-infected cells, and cells subjected to short-term BmLV exposure were infected with recombinant BmNPV expressing luciferase, and BmNPV genome abundance and luciferase activity were measured at the indicated time points. Firefly luciferase (Luc) is a widely used reporter for quantifying gene expression and promoter activity. In this study, the Luc reporter was incorporated into the BmNPV construct and placed under the control of the baculovirus polyhedrin promoter. Luc expression therefore provided a quantitative measure of BmNPV gene expression during infection, without being required for viral replication or serving as a functional viral or host-response factor.

We examined BmNPV genome abundance in the culture supernatant. BmNPV genome abundance was significantly higher in persistently BmLV-infected cells at 72 h post-infection, whereas the difference was no longer significant at 96 hpi (Fig. 1B). Consistently, persistent BmLV-infected cells showed significantly higher luciferase activity than BmLV-free cells following BmNPV infection at 96 hpi (Fig. 1C). Luciferase activity in the persistently BmLV-exposed group was also higher than that in simultaneously BmLV-and BmNPV-coinfected cells, although this difference was not statistically significant.

**Fig. 1.**
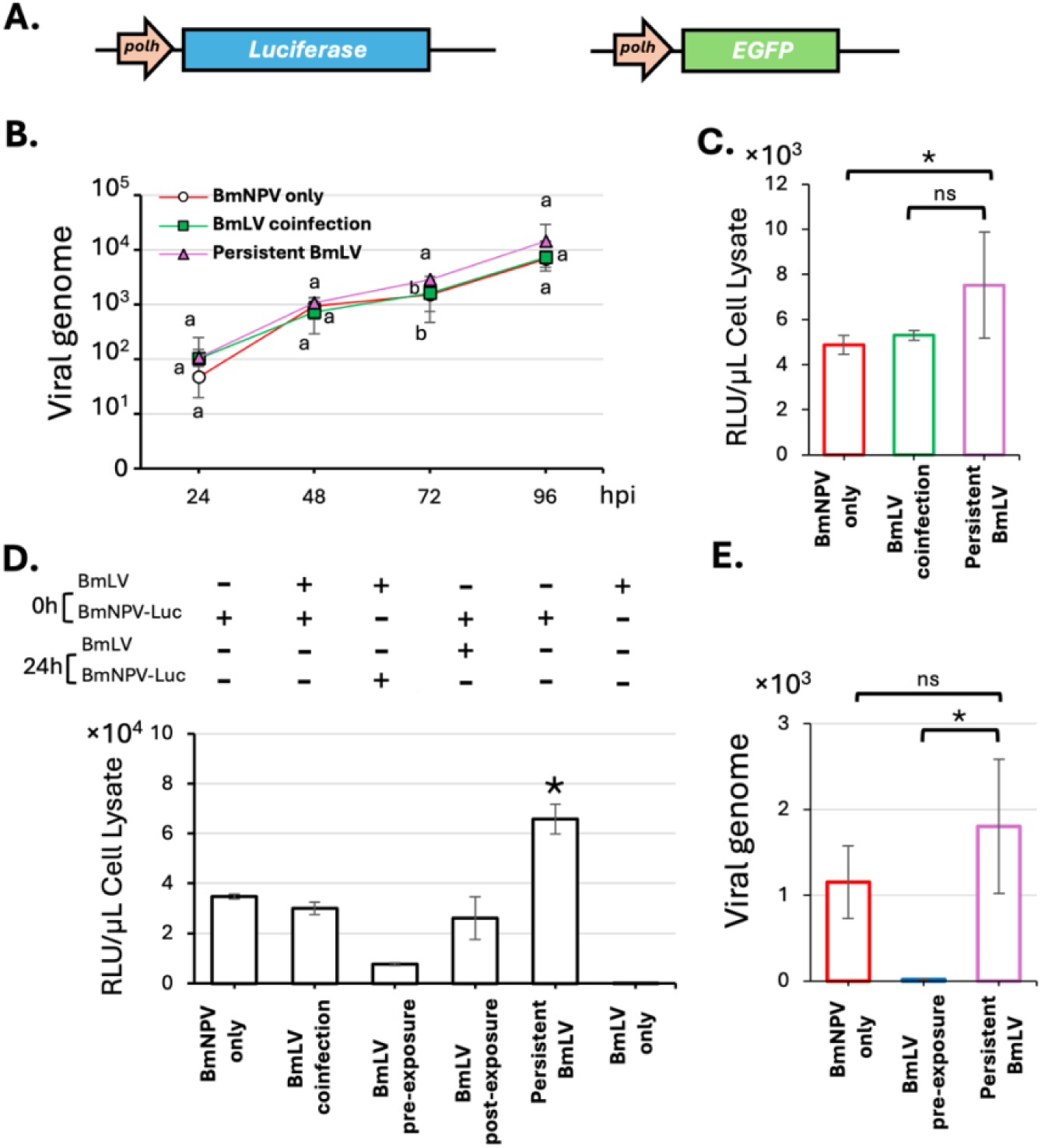
Persistent *Bombyx mori* latent virus (BmLV) infection, but not transient exposure, enhances BmNPV-mediated reporter expression in cultured cells. (**A**) Schematic representations of the recombinant baculovirus constructs encoding the indicated reporter genes. The constructs included firefly luciferase and enhanced green fluorescent protein (EGFP), both expressed under the control of the polyhedrin (*polh*) promoter. Construct diagrams are not drawn to scale. (**B**) Quantification of viral genome accumulation following infection with BmNPV expressing EGFP at 96 h post-BmNPV infection (hpi). BmMAX7050np cells were seeded at 5 × 10^5 cells/well in 12-well plates and infected with BmNPV at an MOI of 0.1. Culture supernatants were collected at the indicated time points, and viral genome abundance was determined by quantitative PCR (qPCR). (**C**) Luciferase activity in BmMAX7050np cells following infection with recombinant BmNPV expressing luciferase at a multiplicity of infection (MOI) of 1 at 96 hpi. (**D**) Luciferase activity in BmMAX7050np cells under the indicated BmLV exposure conditions. Samples were collected at 96 hpi. (**E**) Quantification of BmNPV genome accumulation in the BmLV pre-exposure group at 48 hpi. Cells were infected with BmNPV-Luc at an MOI of 0.1. Data are presented as mean ± standard deviation from at least three independent samples. Luciferase activity data were log10-transformed before statistical analysis. Comparisons between two groups were performed using Welch’s *t*-test, and comparisons among three or more groups were performed using Welch’s one-way ANOVA followed by the Games–Howell post hoc test. Different letters or an asterisk (*) indicate significant differences (*P* < 0.05), whereas the same letters or “ns” indicate no significant difference.

Because the increase in luciferase activity was observed specifically in persistently BmLV-infected groups, we hypothesized that the effects of prior BmLV exposure may depend on persistent infection. We therefore examined whether short-term BmLV exposure before BmNPV infection was sufficient to reproduce the increased reporter protein expression observed in persistently BmLV-infected cells. BmLV was introduced 24 h before BmNPV infection, and this pre-exposure group was compared with BmNPV infection alone, simultaneous BmLV and BmNPV exposure (coinfection), and BmLV exposure 24 h after BmNPV infection (post-exposure) (Fig. 1D). None of the short-term BmLV exposure conditions significantly increased luciferase activity relative to the BmNPV-only group. Consistent with Fig. 1C, persistently BmLV-infected cells showed higher viral genome abundance than pre-exposed cells (Fig. 1E).

Together, these results indicate persistent BmLV infection, but not short-term exposure, enhances BmNPV abundance and BmNPV-mediated reporter expression in cultured cells.

### 3.2 BmLV pre-exposure enhances BmNPV-mediated reporter expression in silkworm larvae

The distinct effects of persistent and short-term BmLV exposure in cultured cells prompted us to examine whether the temporal relationship between BmLV and BmNPV exposure affects BmNPV infection *in vivo*. We established a BmLV pre-exposure model (BmLV exposure 24 h before BmNPV infection) in fifth-instar larvae and compared BmNPV infection alone and simultaneous BmLV and BmNPV exposure (BmLV coinfection) (Fig. 2A).

**Fig. 2.**
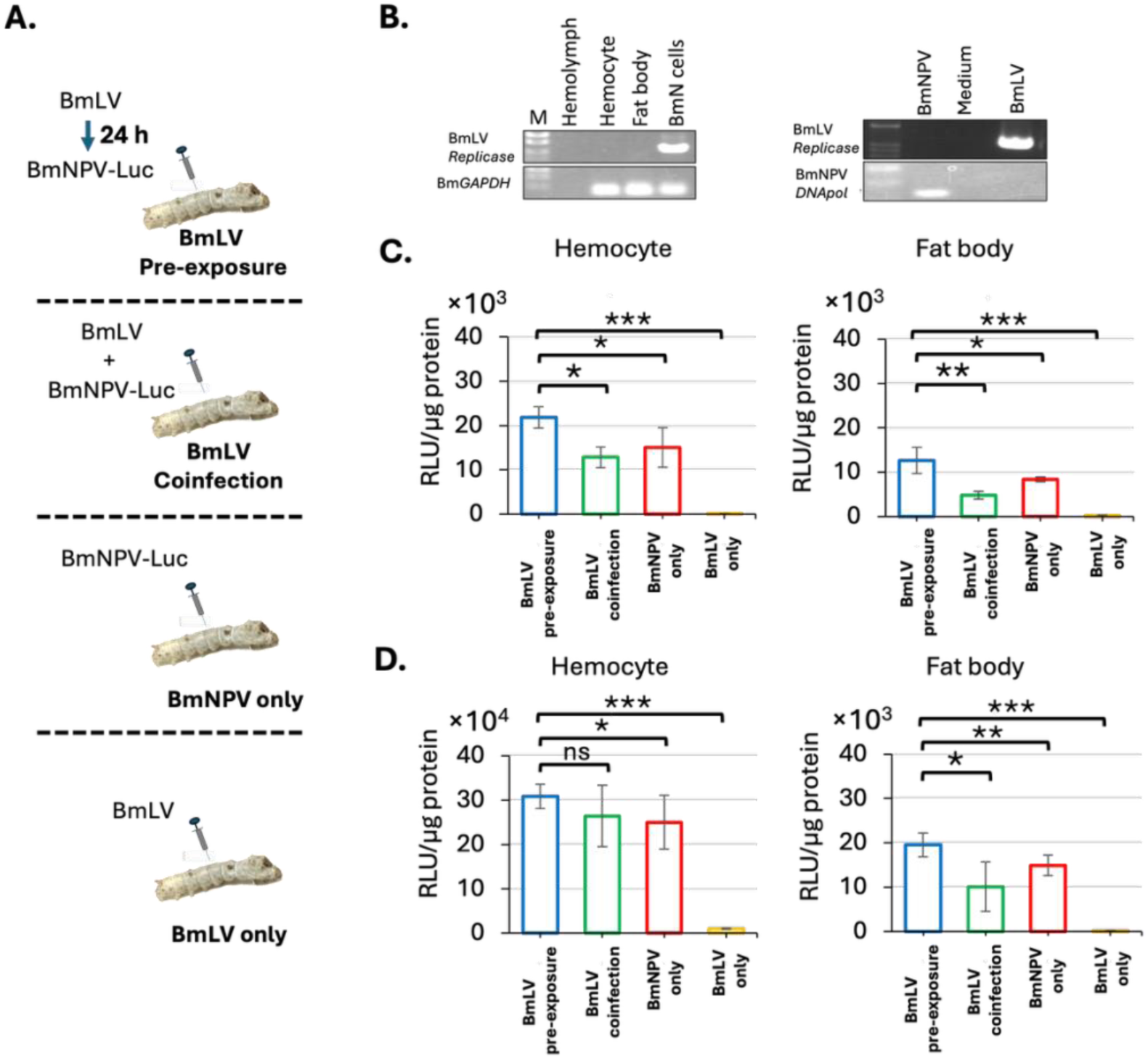
Prior BmLV exposure enhances BmNPV-mediated reporter expression in silkworm larvae. (**A**) Schematic representation of the experimental groups and treatment schedules. (**B**) Assessment of BmLV RNA in tissues from fifth-instar silkworm (*B. mori*) n22 larvae (*n* = 3). Tissues were pooled, total RNA was extracted, and BmLV *replicase* RNA was analyzed by RT-PCR (left panel). *BmGAPDH* was analyzed as an internal control. BmNPV and BmLV inocula and the corresponding control medium were screened for potential cross-contamination (right panel). (**C**) Luciferase activity in n22 fifth-instar larvae following infection with 1 × 10^4 PFU of recombinant BmNPV expressing luciferase (BmNPV-Luc). BmLV exposure was performed according to the schedules shown in panel B. Hemocytes and fat bodies were collected at 96 h post-BmNPV infection (hpi). (**D**) Luciferase activity in f38 fifth-instar larvae under the same experimental conditions. Data are presented as mean ± standard deviation from at least three independent samples. Luciferase activity data were log10-transformed before statistical analysis. Comparisons between two groups were performed using Welch’s *t*-test, and comparisons among three groups were performed using Welch’s one-way ANOVA followed by the Games–Howell post hoc test. An asterisk (*, **, ***) indicates significant differences (*P* < 0.05, *P* < 0.01, *P* < 0.005), respectively, whereas “ns” indicates no significant difference.

Before the larval tissue analysis, we confirmed the BmLV and BmNPV status of the inocula and experimental larvae. Endogenous BmLV RNA was not detected in the larvae before inoculation, whereas BmLV RNA was detected in persistently infected BmN cells used as a positive control (Fig. 2B, left panel). BmLV RNA was not detected in the control medium or in the BmNPV inoculum prepared from BmLV-free BmMAX7050np cells. In contrast, BmLV RNA was detected in the BmLV inoculum, while BmNPV DNA was not detected in this preparation (Fig. 2B, right panel). These results indicated no detectable cross-contamination between the BmLV and BmNPV inocula. No mortality was observed in either f38 or n22 larvae during the five-day observation period following BmLV exposure (Table S3). We selected n22 and f38 because both strains were previously identified as high-expression performers in a comparative screen of inbred silkworm strains for recombinant SARS-CoV-2 spike protein production (Masuda et al., 2022), indicating that they support robust baculovirus-mediated protein expression.

At 48 hpi, no significant difference in luciferase activity was observed between the BmLV-pre-exposed and control groups in n22 larvae (Fig. S1). At 96 hpi, BmLV pre-exposure resulted in significantly higher luciferase activity in hemocytes and fat body than in BmNPV alone or BmLV coinfection groups (Fig. 2C). We next examined whether the effect of BmLV pre-exposure was reproducible in a second genetic background. In f38 larvae, BmLV pre-exposure significantly increased luciferase activity in both hemocytes and fat body compared with BmNPV infection alone at 96 hpi (Fig. 2D). Luciferase activity was also higher in BmLV-pre-exposed larvae than in the coinfection group in the fat body. However, no significant difference was observed between the two groups in hemocytes.

These results demonstrated that exposure to BmLV before BmNPV infection consistently enhanced BmNPV-mediated reporter expression in two *B. mori* genetic backgrounds, although the magnitude of the effect differed between strains.

### 3.3 BmLV pre-exposure increases BmNPV propagation in silkworm larvae

Increase in BmNPV-mediated reporter protein expression following BmLV pre-exposure could result from increased viral propagation. We therefore quantified BmNPV genome abundance following larval infection in fifth instar n22 larvae. BmLV pre-exposed larvae exhibited significantly higher BmNPV genome levels in cell-free hemolymph than BmNPV-only controls at 48 h post-BmNPV infection (Fig. 3A). This increase in BmNPV genome abundance in BmLV pre-exposed larvae remained evident through the final time point examined, 96 hpi.

**Fig. 3.**
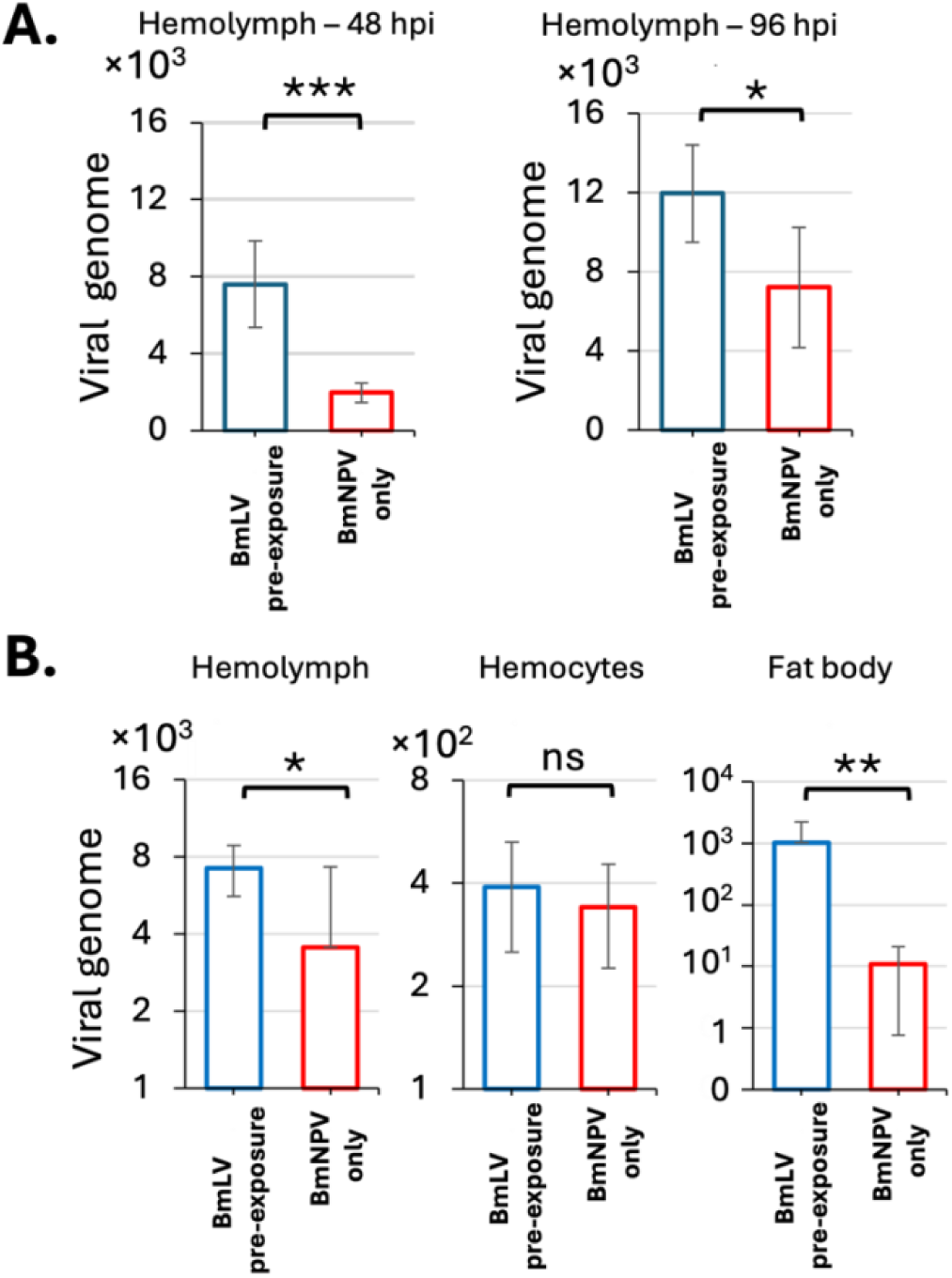
Prior BmLV exposure increases BmNPV propagation in silkworm larvae. (**A**) Absolute quantification of BmNPV genome accumulation in cell-free hemolymph of fifth-instar n22 larvae following infection with 1 × 10^4 PFU of recombinant BmNPV. Samples were collected at 48 and 96 h post-BmNPV infection (hpi), and viral genome abundance was determined by quantitative PCR (qPCR). Viral genome in hemolymph was normalized to virus copies per µL. (**B**) Analysis of BmNPV genome abundance in the f38 larval strain at 96 hpi. Abundance of viral genome in hemocyte and fat body tissues was normalized to *BmGAPDH* abundance. For the BmLV pre-exposure group, BmLV inoculum was administered 24 h before BmNPV infection. The BmNPV-only group received recombinant BmNPV inoculum. Data are presented as mean ± standard deviation from at least four independent samples. Comparisons between experimental groups were performed using Welch’s *t*-test. An asterisk (*, **, ***) indicates significant differences (*P* < 0.05, *P* < 0.01, *P* < 0.005), respectively, whereas “ns” indicates no significant difference.

In the f38 larval strain, we measured viral genome abundance at 96 hpi in hemolymph, hemocytes, and fat body (Fig. 3B). Viral genome abundance was significantly higher in BmLV-pre-exposed larvae in hemolymph and fat body than in BmNPV-only controls, whereas hemocytes showed no significant difference.

These results demonstrate that BmLV pre-exposure increased BmNPV genome accumulation in larvae.

### 3.4 BmLV antigenome RNA is detected in multiple larval tissues during BmLV pre-exposure

Previous studies have reported persistent BmLV infection in silkworm-derived cultured cells and detection of BmLV RNA in the fat body, midgut, and gonads of silkworm larvae following hemocoelic inoculation (Katsuma et al., 2005; Iwanaga et al., 2012; Tsukui et al., 2019; Feng et al., 2021). However, whether BmLV actively replicates following experimental exposure of larvae remains unclear. The enhanced BmNPV genome accumulation observed following BmLV pre-exposure raised the possibility that BmLV may replicate during the pre-exposure period. We therefore examined the tissue distribution of BmLV antigenome RNA after experimental exposure and determined whether this replication-associated RNA remained detectable during subsequent BmNPV infection. To distinguish newly synthesized negative-sense antigenome RNA from input positive-sense genomic RNA, strand-specific RT-PCR was used to detect the BmLV antigenome. The experimental workflow is shown in Fig. 4A.

**Fig. 4.**
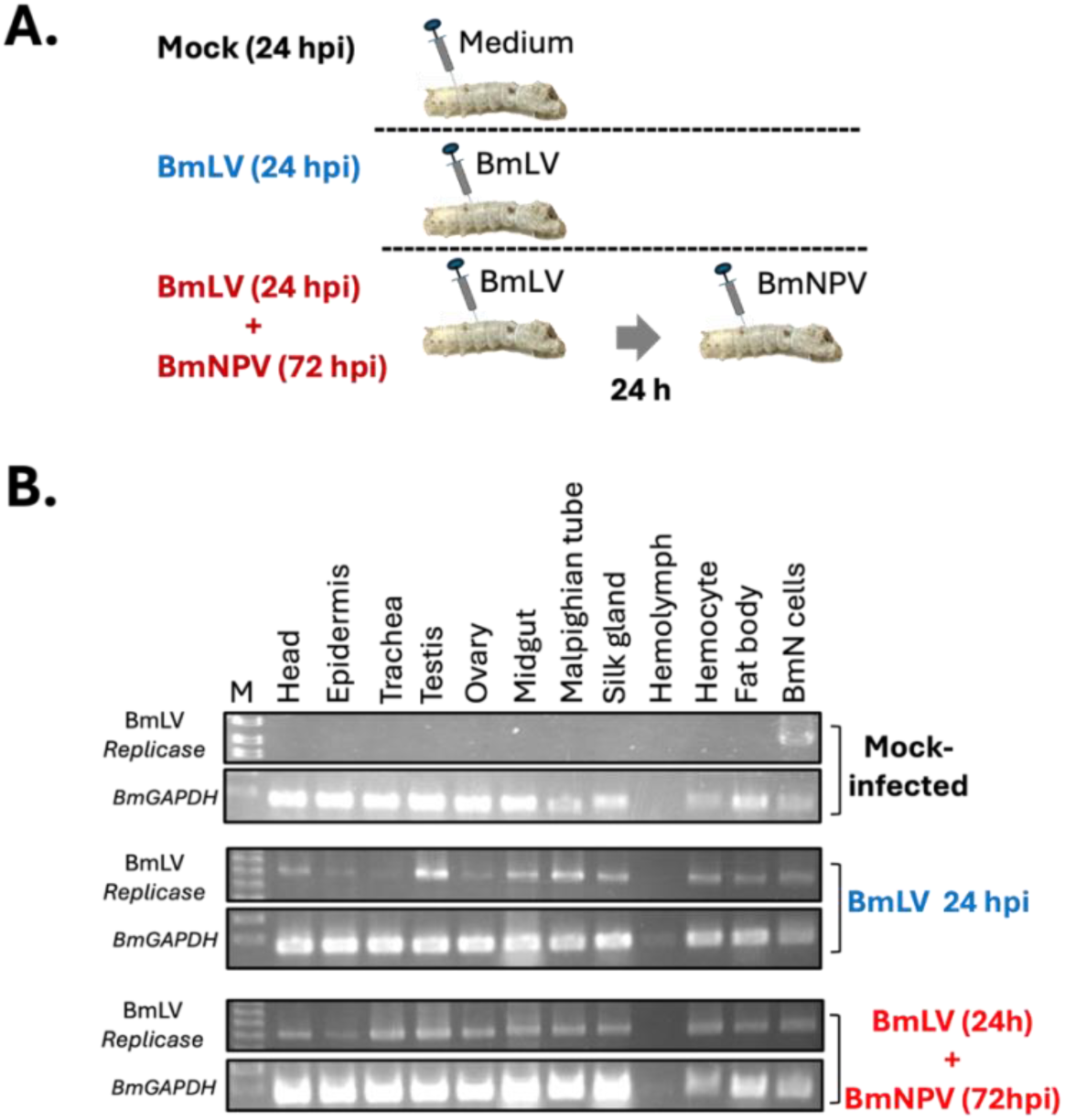
*Bombyx mori* latent virus (BmLV) replicates in multiple larval tissues during larval pre-exposure and remains detectable after subsequent *B. mori* Nucleopolyhedrovirus (BmNPV) infection. (**A**) Experimental groups, infection schedules, and sample collection time points for BmLV and BmNPV inoculation in fifth-instar silkworm (*B. mori)* f38 larvae. BmLV was injected at −24 h; BmNPV at 0 h; sampling at 72 h after BmNPV infection. (**B**) Analysis of BmLV replication by strand-specific RT-PCR. *BmGAPDH* was used as an internal control. Mock-infected groups received control medium (upper panel). BmLV 24 h groups received BmLV inoculum only, and tissues were collected 24 h after BmLV inoculation (middle panel). For the BmLV 24-h + BmNPV 72-h group, larvae received BmLV inoculum followed by BmNPV infection 24 h later, and tissues were collected 72 h after BmNPV infection (lower panel).

No BmLV antigenome RNA was detected in mock-infected larvae in any of the tissues examined (Fig. 4B, upper panel). At 24 h after BmLV inoculation, BmLV antigenome RNA was detected in the midgut, silk gland, hemocytes, fat body, and testis, with weaker signals in the epidermis, ovary, and trachea (Fig. 4B, middle panel). After the subsequent BmNPV challenge, BmLV antigenome RNA remained detectable at 72 h post-BmNPV infection in the tissues in which it had been detected during the pre-exposure period. BmLV antigenome RNA in the epidermis, ovary, and trachea became more pronounced (Fig. 4B, lower panel). As expected, no BmLV antigenome RNA was detected in the cell-free hemolymph fraction during both BmLV exposure and subsequent BmNPV infection.

These results provide evidence that BmLV replicated in multiple larval tissues during the pre-exposure period and that BmLV antigenome RNA remained detectable during subsequent BmNPV infection.

### 3.5 BmLV pre-exposure attenuates antiviral gene induction during subsequent BmNPV infection

The increased BmNPV genome accumulation following BmLV pre-exposure could reflect changes in host antiviral responses. We therefore examined selected components of two antiviral pathways implicated in the response to BmNPV infection: the RNA interference (RNAi) pathway and STING-associated NF-κB signaling. We examined *Dicer-2, R2D2*, and *Ago2* as representative components of the small interfering RNA (siRNA) pathway, whereas we examined *STING* and *Relish* as components of STING-associated antiviral signaling and NF-κB activation, respectively (Nagamine and Sako, 2016; Hua et al., 2018; Jiang, 2021). We focused on the fat body and hemocytes because both are directly involved in BmNPV infection and antiviral responses. Following hemocoelic inoculation, BmNPV initially infects hemocytes and subsequently disseminates to tissues including the fat body, where viral replication becomes prominent (Rahman & Gopinathan, 2004; Katsuma et al., 2012). Both tissues also exhibit differential antiviral gene expression following BmNPV infection (Bao et al., 2010), with extensive infection and antiviral responses detected in the fat body (Feng et al., 2023). We therefore examined these tissues in f38 silkworm larvae to determine whether prior BmLV exposure altered antiviral responses at sites relevant to BmNPV infection and dissemination. The experimental groups are shown in Fig. 5A.

**Fig. 5.**
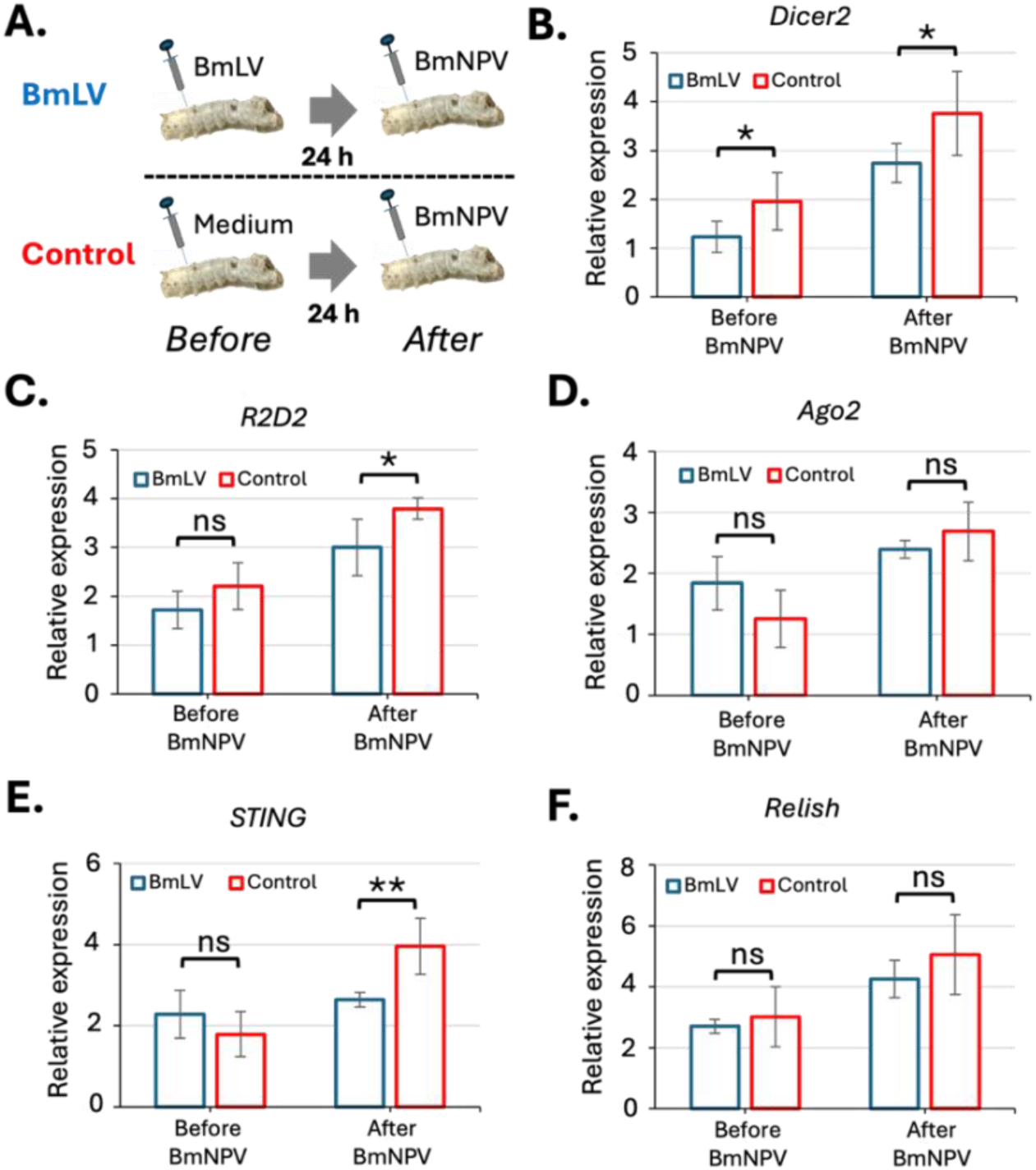
Prior BmLV exposure attenuates antiviral gene induction during subsequent BmNPV infection. (**A**) Experimental design showing the treatment in fifth-instar f38 silkworm larvae. (**B**) Expression of *Dicer-2* in larval fat body following BmLV exposure and subsequent BmNPV infection. BmLV inoculum (20 μL per larva) or control medium was administered, followed by infection with 1 × 10^4 PFU of recombinant BmNPV where indicated. In the “Before BmNPV” condition, samples were collected 24 h after BmLV or control inoculation. In the “After BmNPV” condition, samples were collected 48 h after subsequent BmNPV infection. Relative gene expression was calculated using the 2^−ΔCq method, with *BmGAPDH* used as the endogenous reference gene. (**C**) Expression of *R2D2*. (**D**) Expression of *Ago2*. (**E**) Expression of *STING*. (**F**) Expression of *Relish*. Data are presented as mean ± standard deviation from four independent samples. Data were log10-transformed before statistical analysis. Comparisons between experimental groups were performed using Welch’s *t*-test. Asterisks (*, **, ***) indicate significant differences (*P* < 0.05, *P* < 0.01, *P* < 0.005), respectively, whereas “ns” indicates no significant difference.

After 24 h of BmLV pre-exposure, *Dicer-2* transcript levels were significantly lower in the fat body of BmLV-pre-exposed larvae than in mock-pre-exposed larvae (Fig. 5B). In contrast, transcript levels of *Ago2*, *R2D2*, Relish, and *STING* did not differ significantly between the two groups (Fig. 5C–F). These results indicate that BmLV pre-exposure did not significantly influence the basal expression of most antiviral genes examined.

At 48 h after BmNPV infection, transcript levels of *Dicer-2*, *R2D2*, *Ago2*, *Relish*, and *STING* increased relative to the prior BmNPV infection state in the fat body. However, the extent of transcript increases varied between BmLV-pre-exposed and mock-pre-exposed larvae. *Dicer-2* and *R2D2* transcript levels were significantly lower in BmLV-pre-exposed larvae than in mock-pre-exposed larvae, whereas *Ago2* transcript levels did not differ significantly between the two groups. *STING* transcript levels were also significantly lower following BmLV pre-exposure, while *Relish* transcript levels did not differ significantly between the groups (Fig. 5B–F). Thus, these results suggest that prior BmLV exposure attenuates the expression of selected antiviral genes in the fat body.

In hemocytes, the effects of BmLV pre-exposure were less pronounced than those observed in the fat body (Fig. S2). After 24 h of BmLV pre-exposure (before BmNPV infection), expression of *Relish* was attenuated compared to the mock-exposed groups, whereas no significantly different expression was observed in *STING*, *Dicer-2*, *R2D2*, and *Ago2*. At 48 h after BmNPV infection, transcript levels of *Dicer-2*, *R2D2*, *Ago2*, and *Relish* did not differ between BmLV-and mock-exposed groups. However, *STING* expression increased in the BmLV pre-exposed group relative to the mock-exposed group in the hemocytes.

Overall, BmLV pre-exposure was associated with attenuated induction of selected antiviral genes during subsequent BmNPV infection, particularly *Dicer-2*, *R2D2*, and *STING* in the fat body. This attenuation coincided with increased BmNPV genome accumulation in BmLV-pre-exposed larvae. Prior BmLV exposure may modify antiviral responses during subsequent BmNPV infection.

## Discussion

This study demonstrates that prior BmLV exposure alters subsequent BmNPV infection and that the outcome depends on both the duration and temporal order of exposure. In cultured cells, persistent BmLV infection enhanced BmNPV genome accumulation and reporter expression, whereas 24-h BmLV pre-exposure did not reproduce this effect. In contrast, 24-h BmLV pre-exposure increased BmNPV genome accumulation and reporter expression in larvae, whereas BmNPV alone or simultaneous BmLV and BmNPV did not produce a comparable effect. Thus, short-term BmLV exposure can alter subsequent BmNPV infection in larvae, but not in cultured cells, indicating that this interaction is strongly influenced by the biological context of the host.

A possible explanation is that BmLV exposure in larvae induces tissue-level or systemic responses that are absent or limited in cultured cells. BmLV RNA has been detected in multiple larval tissues following experimental exposure, and transient BmLV propagation has been associated with activation of the Imd and RNAi pathways (Katsuma et al., 2005; Feng et al., 2021). BmNPV, in turn, initially infects hemocytes and subsequently disseminates to multiple tissues, including the fat body, where infection and antiviral responses are prominent (Rahman & Gopinathan, 2004; Katsuma et al., 2012; Bao et al., 2010; Feng et al., 2023). BmLV pre-exposure may alter the tissue state, which could subsequently modify the response to BmNPV infection. However, whether this tissue-level response directly causes the enhanced BmNPV infection remains to be determined.

A previous study demonstrated that BmLV and BmNPV can interact, as simultaneous inoculation of the two viruses increased BmLV accumulation in larvae (Tsukui et al., 2019). In this study, we found that prior BmLV exposure increased BmNPV genome accumulation. The increase in viral genome abundance was detectable before the later increase in reporter accumulation, indicating that enhanced viral propagation may contribute to the subsequent increase in BmNPV-mediated reporter expression. Together, these findings indicate that the temporal relationship between the two viruses can alter the outcome of their interaction.

In cultured cells, a previous study reported that persistent BmLV infection delays BmNPV propagation (Tsukui et al., 2019), whereas we observed increased BmNPV genome accumulation in persistently BmLV-infected cells. Similarly, Iwanaga et al. (2014) found no discernible effect of persistent BmLV infection on BmNPV-mediated recombinant protein production, while persistently BmLV-infected cells in the present study exhibited higher reporter expression. The differences between our findings and those of previous studies may be attributable to variations in cellular background, viral strains, or other experimental conditions.

In cultured-cell analysis, short-term BmLV exposure did not increase BmNPV genome accumulation or reporter protein expression, indicating that the consequences of BmLV are not determined simply by the presence of viral RNA. Moreover, the antiviral gene expression results suggest that BmLV pre-exposure may alter the host physiological state, thereby influencing the outcome of subsequent BmNPV infection. BmLV exposure alone produced relatively limited changes in basal expression of the genes examined, although *Dicer-2* expression was reduced in the fat body. Following BmNPV infection, induction of *Dicer-2*, *R2D2*, and *STING* was significantly lower in BmLV-pre-exposed larvae in the fat body. RNA interference is a major antiviral defense mechanism in silkworms, and STING-associated signaling contributes to antiviral defense against BmNPV (Jiang, 2021; Hua et al., 2018). The reduced induction of selected antiviral genes in BmLV pre-exposed larvae following subsequent BmNPV infection may be associated with increased BmNPV genome abundance, likely by attenuating the host antiviral response.

BmLV-pre-exposure increased BmNPV genome abundance in the fat body but not in hemocytes. These observations are consistent with the expression patterns of the selected antiviral genes, as antiviral gene induction was attenuated in the fat body of BmLV-pre-exposed larvae. In contrast, only minor differences in antiviral gene expression were detected between the two groups in hemocytes. However, despite no significant attenuation of antiviral gene expression in hemocytes, BmLV-pre-exposure increased BmNPV-mediated reporter protein expression in this tissue. This finding suggests that the effects of prior BmLV exposure on subsequent BmNPV infection outcomes may not be explained solely by the antiviral pathways examined in this study and could involve additional host factors or biological processes. Furthermore, transcript abundance does not directly reflect pathway activity, and these findings do not establish a causal relationship between reduced antiviral gene induction and increased BmNPV propagation. Functional manipulation of these pathways in BmLV-pre-exposed larvae will be required to test this possibility.

Previous studies reported that BmLV does not efficiently replicate in *B. mori* larvae following experimental inoculation, although BmLV propagation can be enhanced during BmNPV coinfection (Iwanaga et al., 2012; Tsukui et al., 2019). In contrast, we detected negative-sense BmLV antigenome RNA in multiple larval tissues 24 h after BmLV exposure, before BmNPV challenge. These findings provide evidence of BmLV replication in larvae without BmNPV coinfection. The observed difference from previous studies may reflect variation in silkworm genetic background, rearing conditions, inoculum preparation or dose, sampling time, or assay sensitivity. Although BmLV exposure did not cause detectable mortality or obvious morphological abnormalities, the detection of antigenome RNA in multiple tissues demonstrates that the exposure was not restricted to a single tissue compartment.

Taken together, our findings identify prior viral exposure as a determinant of subsequent BmNPV infection in *B. mori*. They also have practical implications for baculovirus expression systems, as persistent BmLV carriage may influence BmNPV propagation and recombinant protein expression in BmLV-positive silkworm-derived cell lines. More broadly, these findings demonstrate that infection history can shape the outcome of a subsequent viral infection even in the absence of overt disease. Identifying the host changes established by BmLV exposure and determining how they influence subsequent BmNPV infection will be important for understanding sequential viral infections in silkworms.

## Materials and methods

### Insect cell lines and culture conditions

BmN (*B. mori* ovary-derived cell lines) and BmMAX7050np cells were maintained in IPL-41 medium supplemented with 10% fetal bovine serum (FBS) at 27°C. BmMAX7050np cells, derived from G1 embryos of the p50 and n70 strain background as described previously (Abubakar et al., 2026), were used as the BmLV-free cell line. The absence of detectable BmLV in BmMAX7050np cells was confirmed by reverse transcription polymerase chain reaction (RT-PCR) before use. BmN cells are the parental line of BmN4 cells and were previously shown to be persistently infected with BmLV; we used BmN cells as BmLV-positive cells, as described previously (Abubakar et al., 2026).

### Silkworm strains and rearing conditions

Silkworm (*B. mori*) of the n22 and f38 strains, obtained from the National Bio-Resource Project (NBRP) silkworm, were used for all larval experiments. Larvae were reared on mulberry leaves at 25–27°C and used on day 0 or 1 of the fifth instar. Unless otherwise indicated, 4–6 larvae were used per experimental group. Larvae were randomly assigned to treatment groups before viral inoculation.

### Recombinant baculoviruses and BmLV inoculum

Recombinant BmNPV carrying the indicated reporter was generated as described previously (Abubakar et al., 2026). Briefly, the corresponding transfer vector carrying the target gene was introduced into the BmNPV bacmid (Motohashi et al., 2005). Recombinant bacmid DNA was transfected into BmMAX7050np cells to generate BmLV-free virus stocks. BmNPV stocks at passages P3–P5 were used throughout this study. The recombinant baculoviruses used are listed in Table S1, and the constructs are presented in Figure 1A.

BmLV inoculum was prepared from approximately 90% confluent BmN cultures in T25 flasks persistently infected with BmLV (Katsuma et al., 2005; Iwanaga et al., 2012). Culture supernatants were collected, clarified by centrifugation, and passed through a 0.22-μm membrane filter before use. The same BmLV inoculum preparation procedure was used throughout the study. For larval experiments, 20 μL of BmLV inoculum was administered by hemocoelic injection. BmLV-free culture medium collected from BmMAX7050np cultures was used as the corresponding control.

The viral inocula were examined for BmLV RNA and BmNPV DNA by RT-PCR and PCR, respectively, to assess potential cross-contamination between the BmLV and BmNPV inocula.

### Plaque assay for BmNPV stock titration

BmNPV infectious titers were determined by plaque assay based on previously described baculovirus methods (O’Reilly et al., 1992, 1994).

### BmNPV infection and viral genome quantification

For cultured cells, 1 × 10^6 cells were seeded in six-well plates or 5 × 10^5 cells in 12-well plates and incubated for 24 h before infection. Cells were infected with recombinant BmNPV at the indicated multiplicity of infection (MOI). MOIs of 0.1 and 1 were used for baculovirus susceptibility and reporter expression analyses, respectively. For analysis of larval tissues, fifth-instar n22 and f38 larvae were challenged by hemocoelic injection with 1 × 10^4 PFU of BmNPV per larva. Viral genome accumulation was examined at the indicated time points, including 48 and 96 hours post-infection (hpi).

Infected hemocytes or fat body tissues were resuspended in lysis buffer (50 mM Tris-HCl, pH 7.8; 100 mM NaCl; 1 mM DTT; 0.5% CHAPS) and incubated at room temperature for 15 min. Lysates were collected by centrifugation at 14,000 rpm for 5 min at 4°C, and DNA was purified by the phenol-chloroform method. Approximately 100 ng of DNA was used as a template in qPCR. BmNPV genome abundance was normalized to GAPDH.

For rapid quantification of BmNPV genomes in culture supernatants or cell-free hemolymph, samples were mixed with an equal volume of 0.04% SDS and incubated at 70°C for 10 min to lyse cells and disrupt viral particles (Simoliunas et al., 2021). After cooling to room temperature, 1 μL of the treated sample was used directly as the template for qPCR with KOD SYBR qPCR Mix (TOYOBO, Japan). Absolute BmNPV genome copy numbers were determined using a standard curve generated from a purified PCR amplicon corresponding to the BmNPV DNA polymerase target region. The amplicon was generated with the same qPCR primers, verified by agarose gel electrophoresis, purified with a Favorgen PCR/Gel purification kit (FARVOGENE, Taiwan), and quantified with a NanoDrop One spectrophotometer (Thermo Fisher Scientific, USA). Ten-fold serial dilutions of the purified DNA were used to generate the standard curve. BmNPV genome abundance was quantified using primers targeting the BmNPV DNA polymerase gene (Table S2).

### BmLV infection and exposure models

BmLV-free BmMAX7050np cells were used to establish persistent and short-term BmLV exposure models. For the persistent infection model, cells were exposed to BmLV-containing culture supernatant (from BmN cells) and continuously cultured for 80–100 passages. Persistent BmLV infection was confirmed by RT-PCR detection of BmLV RNA. For analysis of short-term exposure, BmLV inoculum was added to the culture medium at a final concentration of 5% (v/v). Cells were exposed to BmLV for 24 h before or after BmNPV infection, or simultaneously with BmNPV infection. BmLV-free culture medium was used as the control.

To examine whether the timing of BmLV exposure affected subsequent BmNPV infection, three exposure schedules were compared: BmLV exposure 24 h before BmNPV infection (pre-exposure), simultaneous BmLV and BmNPV exposure (coinfection), and BmLV exposure 24 h after BmNPV infection (post-exposure). BmNPV-only and BmLV-only groups were included where indicated.

Fifth-instar n22 and f38 larvae were injected hemocoelically with 20 μL of BmLV inoculum and maintained for 24 h before subsequent BmNPV challenge. This group was designated the BmLV pre-exposure group. Control larvae received 20 μL of BmLV-free culture medium. Pre-exposed larvae were challenged with 1 × 10^4 PFU of BmNPV 24 h after BmLV exposure. For coinfection analysis, BmLV and BmNPV were administered simultaneously.

BmNPV-mediated reporter protein expression was assessed at 48 or 96 hpi, depending on the purpose. BmNPV genome accumulation was assessed at 48 and 96 hpi. For analysis of BmLV replication, tissues were collected 24 h after BmLV exposure and 72 h after subsequent BmNPV infection. For antiviral gene expression analysis, samples were collected 24 h after BmLV exposure and 48 h after subsequent BmNPV infection.

### Larval tissue and hemolymph collection

Hemolymph was collected from individual larvae by puncturing a proleg and allowing the hemolymph to drain into sterile microcentrifuge tubes containing ∼1 mg of 1-phenyl-2-thiourea (Wako, Japan) to prevent melanization. Collected hemolymph was centrifuged at 1,500 rpm for 5 min to separate hemocytes from the cell-free hemolymph fraction. The resulting cell-free hemolymph was collected for viral genome quantification, while the hemocyte pellet was used for subsequent analyses as indicated.

For tissue-specific analyses, larvae were dissected at the indicated time points, and the fat body, midgut, silk gland, epidermis, head, Malpighian tubules, trachea, ovary, and testis were collected in sterile microcentrifuge tubes as required for each experiment. Samples were washed three times with 1× phosphate-buffered saline (PBS) and transferred immediately to the appropriate lysis or extraction solution and stored under conditions appropriate for subsequent reporter expression or PCR analysis.

### Luciferase reporter assay

For luciferase analysis, harvested cells at 96 hpi were resuspended in lysis buffer containing 50 mM Tris-HCl (pH 7.8), 100 mM NaCl, 1 mM DTT, and 0.5% CHAPS and incubated at room temperature for 15 min. Lysates were centrifuged at 14,000 × g for 5 min at 4°C. Thirty microliters of the resulting supernatant were mixed with 70 μL of D-luciferin potassium salt solution (Wako, Japan) in a 96-well plate. Luminescence was measured using a VICTOR Nivo multimode plate reader (PerkinElmer, UK) and expressed as relative luminescence units (RLU). For larval samples, hemocytes and fat bodies were collected at the indicated time points (48 or 96 hpi) and processed using the same protocol. Cultured cell lysates were normalized to cell number, whereas larval samples were normalized to total protein content.

### Protein quantification

Protein concentrations were determined using the DC Protein Assay (Bio-Rad, Hercules, CA, USA) according to the manufacturer’s instructions. The measured protein concentrations were used to normalize the amount of input material for luciferase assays of larval tissue samples.

### Viral RNA extraction

Total RNA was extracted from cultured cells and larval tissues using ISOGEN II (NIPPON GENE, Japan) according to the manufacturer’s instructions. RNA concentration was determined using a NanoDrop One spectrophotometer (Thermo Fisher Scientific, USA). For viral RNA detection, reverse transcription was performed with 2 µg total RNA using random primers or gene-specific primers in a total reaction volume of 20 µL.

### Reverse transcription PCR (RT-PCR) and quantitative PCR

Conventional and quantitative PCR were performed using KOD One PCR Master Mix and KOD SYBR qPCR Mix (TOYOBO, Japan), respectively. PCR cycling conditions were performed according to the manufacturers’ protocols, with primer annealing temperatures set at approximately 3–5 °C below the calculated melting temperature of each primer pair. In qPCR analysis, 40 cycles were performed, whereas in conventional RT-PCR, 32–40 cycles were performed, depending on the target. Primer sequences are listed in Table S2. BmNPV genome accumulation was quantified using primers targeting the BmNPV DNA polymerase gene, whereas BmLV genomic RNA was detected using BmLV-specific primers targeting the viral replicase region. For gene expression analysis, 2 µg total RNA was treated with DNase I (Roche, Switzerland), and cDNA was synthesized from total RNA using ReverTra Ace (TOYOBO, Japan) with oligo(dT) primers in a total reaction volume of 20 µL. Reverse transcription was performed at 42 °C for 90 min, and enzyme inactivation was performed at 99 °C for 5 min for reactions using oligo(dT) primers. For reactions using random primers, an additional incubation at 30 °C for 10 min was performed before the reverse transcription steps.

To assess BmLV replication, strand-specific reverse transcription was performed using a BmLV-specific primer complementary to the negative-sense BmLV antigenome. The resulting cDNA was subjected to PCR using BmLV replicase-specific primers. RNA samples were treated with DNase I before reverse transcription to exclude DNA contamination. Detection of the negative-sense antigenome was interpreted as evidence of BmLV replication-associated RNA synthesis.

Relative gene expression was calculated using the 2^−ΔCq method, with *GAPDH* used as the internal reference gene.

### Antiviral gene expression analysis

Expression of selected antiviral genes was analyzed by RT-qPCR. The genes examined were *B. mori Dicer-2*, *R2D2*, *Ago2*, *Relish*, and *STING*, which represent key components of the RNA interference and STING-dependent antiviral pathways involved in the host response to BmNPV infection (Nagamine and Sako, 2016; Hua et al., 2018; Jiang, 2021). *Dicer-2*, *R2D2*, and *Ago2* were selected to assess the RNAi response, whereas *Relish* and *STING* were selected to assess the STING-dependent antiviral signaling pathway.

For analysis of the antiviral host response to BmLV exposure in larvae, samples were collected at 24 h after BmLV inoculation. For sequential infection analysis in larvae, samples were collected 48 h after BmNPV infection, corresponding to 72 h after the initial BmLV exposure. Fat body and hemocyte samples were analyzed separately. Relative transcript expression was calculated using the 2^−ΔCq method, with *GAPDH* used as the internal reference gene.

## Statistical analyses

Statistical analyses were performed using Microsoft Excel and Google Colaboratory. Comparisons between two groups were performed using Welch’s *t*-test. Comparisons among three or more groups were performed using Welch’s one-way ANOVA followed by the Games–Howell post hoc test. Where appropriate, BmNPV genome abundance and reporter expression data were log10-transformed before statistical analysis. Relative gene expression was calculated using the 2^−ΔCq method, with *GAPDH* used as the internal reference gene. Data are presented as mean ± standard deviation (SD) from 3–6 replicates, as indicated in the corresponding figure legends. A *P* value < 0.05 was considered statistically significant.

## CRediT authorship contribution statement

Conceptualization – Nura Abubakar, Takahiro Kusakabe, Masato Hino

Writing – original draft – Nura Abubakar

Writing – review & editing – Masato Hino, Hiroaki Mon, Tsuguru Fujii, Toshiaki Fujimoto, Takahiro Kusakabe

Validation – Takahiro Kusakabe, Masato Hino

Supervision – Masato Hino, Takahiro Kusakabe

Project administration – Nura Abubakar, Masato Hino

Methodology – Nura Abubakar, Masato Hino, Takahiro Kusakabe, Hiroaki Mon, Tsuguru Fujii, Toshiaki Fujimoto

Investigation – Nura Abubakar, Masato Hino, Takahiro Kusakabe, Hiroaki Mon

Funding – Takahiro Kusakabe, Masato Hino

Formal analysis – Nura Abubakar

Data curation – Nura Abubakar

Software and visualization – Nura Abubakar

## Acknowledgement

This work was supported by JSPS KAKENHI Grant Number 26K18066.

## Appendix A. Supporting information/Supplementary material

Supplementary materials (Fig. S1 & S2, Table S1, S2, & S3)

### Data availability

All data generated or analyzed during this study are included in this article and its supplementary information files.

