## Supplementary material for "Prior *Bombyx mori* latent virus exposure alters subsequent BmNPV infection in the silkworm, *B. mori*": Table S1, Table S2, Table S3, Fig. S1, Fig. S2

### Supplementary materials

#### Table S1: Recombinant baculoviruses used in this study

The virus strains, corresponding baculovirus backbones, and target genes are listed. These recombinant viruses were used to evaluate viral propagation and reporter expression.

| Virus name | BmNPV strain | Expressing protein | Reference |
|---|---|---|---|
| BmNPV- Luc | Q4 | Luciferase | Abubakar et al., 2026 |
| BmNPV-EGFP | T3 | EGFP | Abubakar et al., 2026 |

**Abbreviations:** *B. mori* nucleopolyhedrovirus (BmNPV); luciferase (Luc); enhanced green fluorescent protein (EGFP).

**Table S2:** Primers used in this study for RT-PCR, RT-qPCR, and strand-specific RT-PCR analyses

Primer sequences and corresponding GenBank accession numbers are listed. BGAPDH_F and B_GAPDH_R (Abubakar et al., 2026) amplify cDNA to produce a 417 bp product and were used for conventional RT-PCR, whereas BGAPDH_F and B_GAPDH_R2 amplify a 157-bp product and were used as the reference gene for RT-qPCR. BmLV_F and BmLV_R (Abubakar et al., 2026) amplify a 2,090-bp BmLV product, whereas BmLV_F and qBmLV_R amplify a 1,054-bp product. The BmLV-strand primer was used for strand-specific reverse transcription to detect BmLV antigenome RNA. Oligo(dT), random 9-mer, and strand-specific primers were used for reverse transcription, as indicated.

| Target | Primer | Oligo nucleotide sequence (5’------3’) | GenBank Accession No. |
|---|---|---|---|
| *BmNPV DNApol* | DNApol_F<br>DNApol_R | GATTACGGTGGCTCGGCATT<br>CGAACAAACGCAACGGGTAC | D16231.1 |
| BmLV *Replicase* | BmLV_F<br>BmLV_R<br>qBmLV_R | CTTTCCAAATGTTGCGACACTAAGCGGCCC<br>CTCGGAAG<br>CGCTGAGGGTTTGGGCGCCATTCTCATAAG<br>GGAGATC<br>GATGACAAATTGGAGGGCTG | NC_038331 |
| *BmDicer-2* | Dicer-2_F<br>Dicer-2_R | ATGATGACGACGAGGTTCCC<br>TCGTGTGGTAAAATCGAGCA | XM_0213521<br>83.3 |
| *BmR2D2* | R2D2_F<br>R2D22_R | CTGTCTCTGCTTCTGCACGA<br>CAGTAGCAAATGGTGGGGGT | AB566385.1 |
| *BmAgo2* | Ago2_F<br>Ago2_R | CTCCGCTGCCAAAAATCACC<br>ACCCGTTGCCATATGCTTCA | NM_0010435<br>30.2 |
| *BmSTING* | STING_F<br>STING_R | AGCCTTTATAATTGACGGAGCA<br>ACCATGATCTTCTTAGGCTGTGT | XM_0049265<br>81.5 |
| *BmRelish* | Rel_F<br>Rel_R | TGAAATGCCGAAAACTGAGC<br>CACTCGGGCTCCACATCG | NM_0011024<br>65.1 |
| *BmGAPDH* | BGAPDH_F<br>BGAPDH_R<br>BGAPDH_R2 | ACAAACTGTCTTGCCCCACT<br>AGAGTGTGAATCACCAATGAAG<br>TGTTTTGTTGAGCACCACGG | EU141495 |
| RT | BmLV-Strand | TAGCATGGCTTTCACCAATTTA | NC_038331 |
| RT | Oligo(dT) | TTTTTTTTT | NA |
| RT | Random 9-mers | NNNNNNNNN | NA |

NA – not applicable; RT – reverse transcription. GAPDH – Glyceraldehyde-3-phosphate dehydrogenase (Internal reference control). Forward (_F) and reverse (_R) primers.

**Table S3:** Survival of silkworm larvae following BmLV exposure *B. mori* strain, treatment, number of larvae (n), and mortality are shown. Larvae were injected with 20 µL of either BmLV or control inoculum (medium) and monitored daily for five consecutive days. No mortality was observed during the observation period. f38 and n22 are distinct silkworm strains used for this analysis.

| **Strain** | **Treatment** | ***n*** | **Mortality *n*, (%)** |
|---|---|---|---|
| f38 | BmLV | 12 | 0 (0) |
| | Control | 12 | 0 (0) |
| n22 | BmLV | 12 | 0 (0) |
| | Control | 12 | 0 (0) |

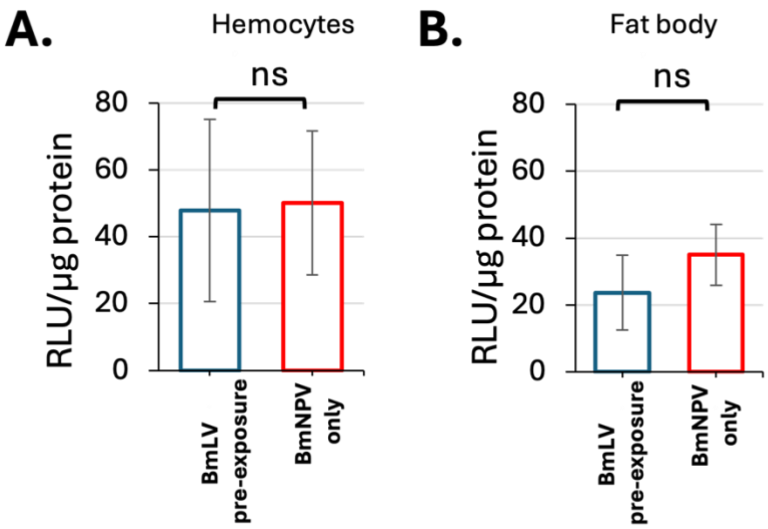


**Fig. S1**. BmNPV-mediated reporter protein expression did not differ significantly between BmLV-pre-exposed larvae and control groups. Luciferase activity was analyzed at 48 hpi in fifth-instar n22 larvae following infection with 1 × 10^4 PFU of recombinant BmNPV expressing luciferase (BmNPV-Luc). BmLV was administered 24 h before BmNPV infection in BmLV-pre-exposed larvae, whereas control groups received BmNPV alone.

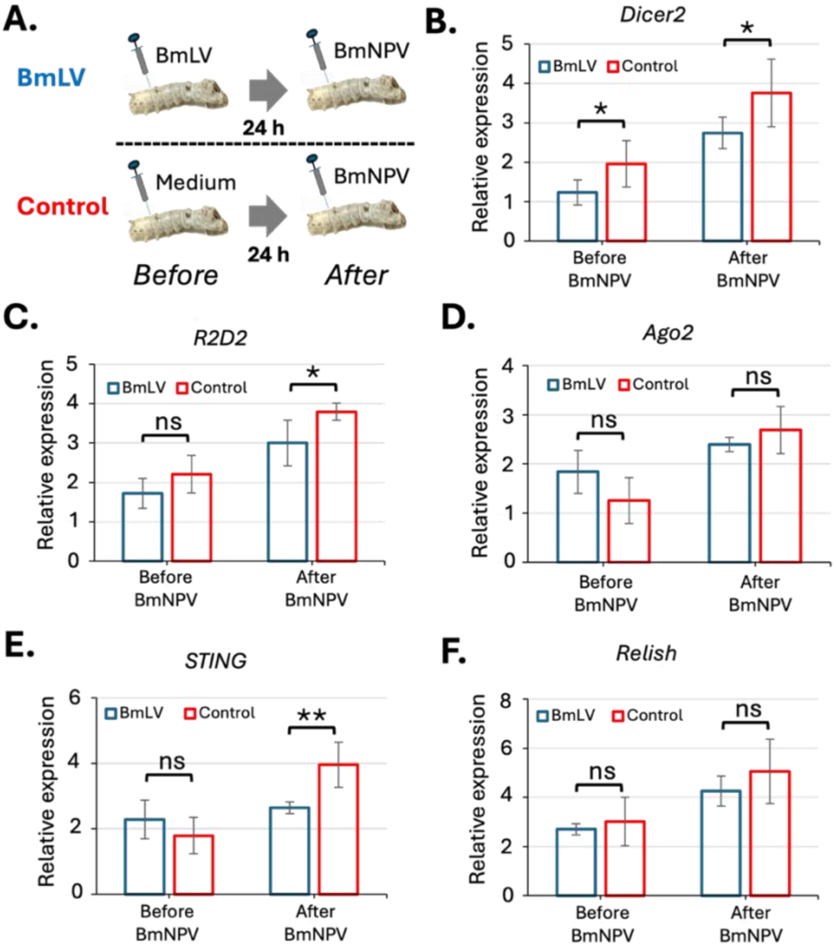


**Fig. S2.** Prior BmLV exposure has limited effect on antiviral gene induction in hemocytes during subsequent BmNPV infection. (**A**) Experimental design showing the treatment and sample collection schedules in fifth instar f38 silkworm larvae. (**B**) Expression of *Dicer-2* in larval hemocytes following BmLV exposure and subsequent BmNPV infection. BmLV inoculum (20 μL per larva) or control medium was administered, followed by infection with 1 × 10^4 PFU of recombinant BmNPV where indicated. In the "Before BmNPV" condition, samples were collected 24 h after BmLV or control inoculation. In the "After BmNPV" condition, samples were collected 48 h after subsequent BmNPV infection. Relative gene expression was calculated using the 2^−ΔCq method, with *BmGAPDH* used as the endogenous reference gene. (**C**) Expression of *R2D2*. (**D**) Expression of *Ago2*. (**E**) Expression of *STING*. (**F**) Expression of *Relish*. Data are presented as mean ± standard deviation from four independent samples. Data were log10-transformed before statistical analysis. Comparisons between experimental groups were performed using Welch's *t*-test. Asterisks (*, **, ***) indicate significant differences ($P < 0.05$, $P < 0.01$, $P < 0.005$), respectively, whereas "ns" indicates no significant difference.
